# Emergent blink rate in early childhood is associated with neural origins of executive function

**DOI:** 10.64898/2026.02.02.703245

**Authors:** Ryuta Kuwamizu, Nozomi Yamamoto, Kota Otani, Yusuke Moriguchi

## Abstract

Executive function develops rapidly in early childhood and predicts later success in life, yet its neurobiological correlates during this period remain poorly understood. In adults, executive function is supported by prefrontal–striatal circuits under neuromodulatory control, including dopamine. Here, we show that spontaneous eye-blink rate (sEBR), a physiological output influenced by neuromodulatory systems, is associated with the developmental trajectory of executive function and prefrontal functional differentiation. We used fNIRS to monitor prefrontal cortex (PFC) activation in 113 children (35–80 months) during a cognitive flexibility task. With increasing age, the children’s task performance improved, and their PFC activation showed greater right-hemisphere and dorsal bias, reflecting functional differentiation. Importantly, higher sEBR was also associated with this right-dorsal PFC activation and with better task performance. The current findings demonstrate that emergent sEBR is associated with the neural origins of executive function development in young children. This highlights the potential of sEBR to serve as a unique and previously unrecognised indicator of this critical developmental neural mechanism.

## Introduction

Executive function—the higher-order cognitive processes enabling goal-directed behaviour and self-control—is essential to quality of life and is an important factor predicting academic achievement, social adjustment, and health across the lifespan^1–3^. Executive function starts to develop during infancy, showing rapid changes during preschool years, and continues to develop through adolescence and into adulthood^4,5^. This trajectory is thought to reflect the functional maturation of the prefrontal cortex (PFC) and associated neural networks^6–8^. According to a prevailing working hypothesis, task-related PFC activation shifts with age—from diffuse, widespread engagement in early childhood to more focal, specialised configurations that rely on efficient networks with other brain regions in later development^9–11^. This transition signifies a functional differentiation of neural recruitment, marking the refinement of prefrontal organization. However, it remains largely unknown whether this functional reorganization already begins to emerge during the preschool years and what neural mechanisms might modulate it^12^.

In adults, executive function is controlled by neuromodulatory systems, including dopaminergic tone. Accumulating evidence demonstrates that the dopaminergic transmission originating in the midbrain is linked with executive function via function in the dorsolateral PFC (DLPFC) and striatum, which is to say the frontostriatal networks^13,14^. Dopamine projections to the PFC are thought to enhance the signal-to-noise ratio of neural representations^15^, stabilizing goal-relevant information, whereas dopaminergic input to the striatum acts as a gate that enables selective updating of task rules^16–18^. Given its critical involvement in both cognitive switching and updating in adults, basic maturation in these circuits may also contribute to the rapid emergence of executive function and efficient cortical recruitment during early childhood. Adults exhibit stronger functional connectivity in the midbrain and lateral PFC (LPFC) during switching tasks compared to preadolescents^19^, supporting the importance of the mechanism underlying executive function development.

However, studies focussing on the neuromodulatory systems of young-child development with rapid executive function improvement are non-existent. PET and postmortem human brain studies suggest that the striatal dopaminergic system rapidly matures by adolescence, likely as a result of the rapid development seen in infancy and childhood^20,21^. On the other hand, resting-state fMRI findings indicate that frontostriatal functional connectivity strengthens only after adolescence^22^, suggesting that it is still developing during early childhood. Interestingly, Moriguchi and Shinohara^23^ reported that polymorphisms in the catechol-O-methyltransferase (COMT) gene—a key regulator of prefrontal dopaminergic tone—are associated with switching performance and right DLPFC (rDLPFC) activation in 3- to 6-year-old children. This finding suggests that the dopaminergic system may already play a functional role in the developing prefrontal function during early childhood. Nevertheless, the absence of non-invasive techniques for tracking the development of neuromodulatory systems means that this mechanism has remained largely unexplored.

To address this challenge, we turned to spontaneous eye-blink rate (sEBR), a physiological output influenced by central neuromodulatory and arousal-related systems^24–29^. Blinking is tightly coupled with brainstem arousal regulation and exhibits sensitivity to changes in internal state^30^. Across adult and clinical studies, individual differences in baseline sEBR have been associated with frontostriatal-dependent functions including executive function^25,27,28,31,32^. Infant studies further suggest that sEBR varies with learning contexts linked to dopaminergic function^33–37^. Several animal, PET, and clinical studies have also reported relationships between sEBR and striatal dopaminergic measures, including D2 receptor function^24–26,28,38,39^ and pharmacological modulation^24–26,38,39^, although the neurochemical specificity of sEBR remains debated^26,40–42^. Interestingly, blink rates are exceptionally low in infancy but increase several-fold between approximately 1 and 5 years of age, and substantial inter-individual differences emerge over the same period^43–45^. Together, these observations motivated us to treat sEBR as a scalable trait-like physiological signature linked to neuromodulatory maturation, with dopamine as one contributor, and to test whether it covaries with executive function development and prefrontal functional differentiation in early childhood.

Here, we provide novel evidence linking early changes in spontaneous eye blinking to the rapid development of executive functions and their neural underpinnings in early childhood (35–80 months). Applying fNIRS during a cognitive flexibility task widely used to measure executive function in young children^7,46^ (Fig. 1), we first confirmed developmental trajectories for PFC activity: older children showed superior performance and greater task-dependent functional differentiation in the PFC (i.e., a right-hemisphere and dorsal bias). Critically, we demonstrate that a higher sEBR is associated with both superior cognitive flexibility and more specialised functional activation in the rDLPFC. These findings are the first to suggest that developmental increases in sEBR are associated with executive function and PFC functional maturation, offering insight into the neural origins of early executive function development.

**Fig. 1.**
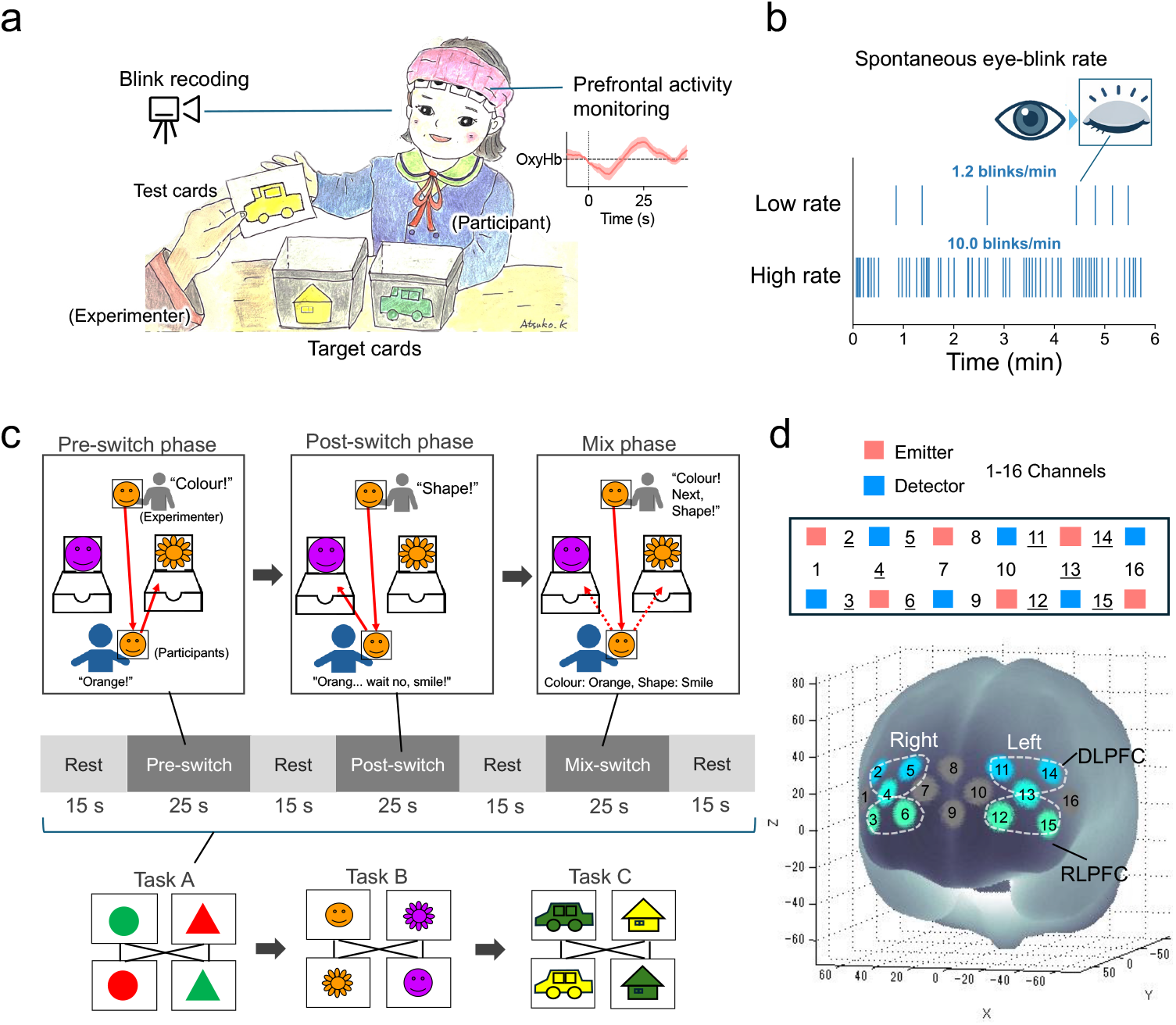
Experimental procedures. **(a)** Schematic illustration of the experimental settings. Participants performed the Dimensional Change Card Sort (DCCS) task while prefrontal haemodynamics and spontaneous eye-blink rate (sEBR) were recorded. **(b)** Spontaneous eye-blink rate (sEBR). Eye blinks were identified from video recordings to calculate sEBR (blinks/min). Raster plots from two example participants illustrating low vs high sEBR during the 6-min are pictured. **(c)** Design of the DCCS task. Top panel: Schematic of the task. The task consists of three phases: a pre-switch phase (sorting by the first dimension), a post-switch phase (sorting by the second dimension), and a mix-switch phase (alternating between the two dimensions). Middle panel: Block design for fNIRS measurement. A 15-s rest period preceded each 25-s task phase. Bottom panel: The stimulus sets used in the task. **(d)** fNIRS probe placement. The 16 measurement channels mapped onto a standard brain template. The measurement area covered the bilateral dorsolateral prefrontal cortex (DLPFC) and rostrolateral prefrontal cortex (RLPFC).

## Results

### sEBR predicts switching performance

First, we recorded the rate of correct responses during the pre-switch, post-switch, and mix phases of the Dimensional Change Card Sort (DCCS) task. Consistent with previous research^23^, the rate of correct responses significantly decreased from the pre-switch phase to the post-switch and mix phases (Friedman test: χ^2^(2) = 68.82, *P* < 0.001; post-hoc tests: *Ps* < 0.001; Supplementary Fig. 1). Next, the percentage of successful shifting (= switching accuracy) was analysed as primary task performance. As expected, switching accuracy improved with age in months (Spearman’s *rho* = 0.68, *P* < 0.001, Fig. 2a). sEBR also increased with age (*rho* = 0.29, *P* = 0.002, Fig. 2b). Critically, supporting our main hypothesis, sEBR was positively correlated with switching accuracy (*rho* = 0.35, *P* < 0.001, Fig. 2c). This association remained significant after controlling for the effect of age (*rho* = 0.22, *P* = 0.022, Fig. 2c). These results suggest that sEBR predicts the development of cognitive flexibility independent of age. Exploratory and robustness analyses further support these findings and are reported in Supplementary Fig. 1b–d (phase-wise analyses) and Supplementary Note 2 (sex- and preschool-adjusted analyses).

**Fig. 2.**
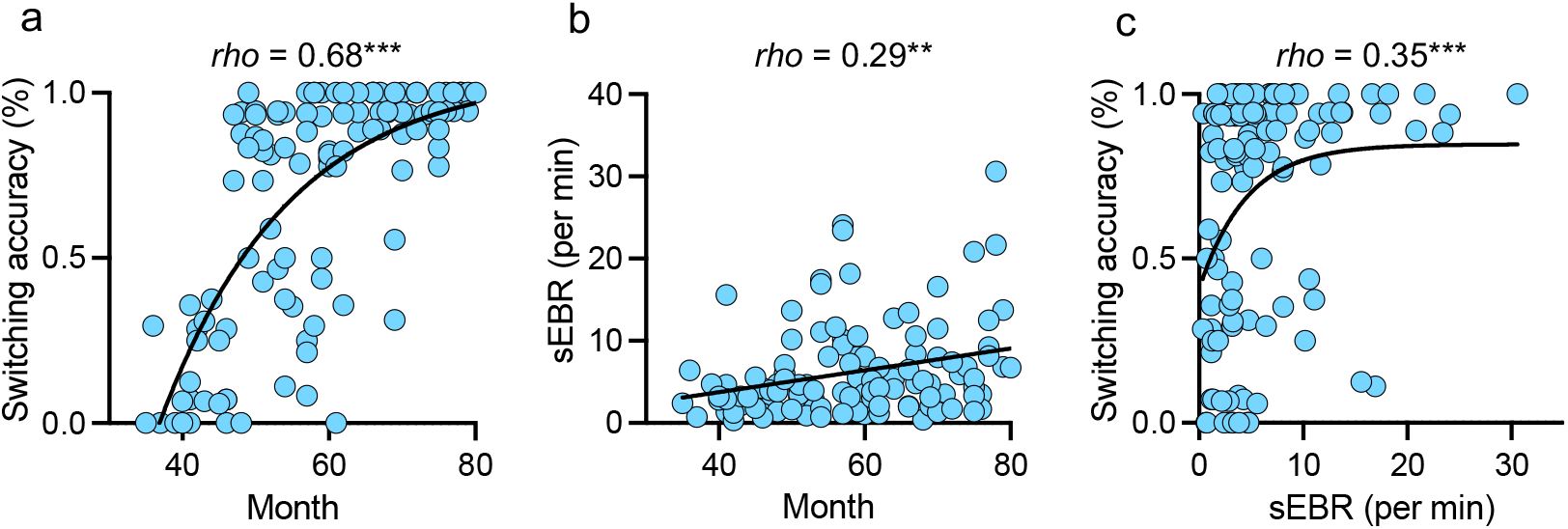
Spontaneous eye-blink rate (sEBR) predicts switching performance. **(a)** Scatterplot of age in months versus switching accuracy. The curve shows a nonlinear fit from an exponential plateau model to capture the ceiling effect. **(b)** Scatterplot of age in months versus sEBR. sEBR increases with age. The line denotes the linear regression fit. **(c)** Scatterplot of sEBR versus switching accuracy. Higher sEBR is associated with greater switching accuracy; this association remains significant after statistically controlling for age. The curve shows a nonlinear fit from an exponential plateau model to capture the ceiling effect. ***P < 0.001, **P < 0.01.

### Cognitive switching induced LPFC activity

To examine task-related prefrontal activation, we fitted a linear mixed-effects model (LMM) to oxy-Hb change with task phase (pre, post, mix), hemisphere (left, right), and region (dorsolateral, rostrolateral) as fixed factors and participant as a random intercept. This model tested the overall effects of the DCCS task on prefrontal activation. The analysis of variance (ANOVA) revealed significant main effects of phase (*F*(2, 1205.2) = 38.44, *P* < 0.001), hemisphere (*F*(1, 1207.2) = 20.07, *P* < 0.001), and region (*F*(1, 1207.2) = 7.14, P = 0.008). No significant interactions were observed in this model (all *Ps* > 0.20). Post-hoc comparisons on estimated marginal means showed that oxy-Hb activity was significantly higher during the switch phase (pre vs. post: *t*(1205) = 7.72, *P* < 0.001; pre vs. mix: *t*(1204) = 7.46, *P* < 0.001) and compared with the pre phase, while no difference was found between the mix and post phases (t(1205) = 0.28, P = 0.958) (Fig. 3a–c). One-sample *t*-tests against baseline further clarified this pattern. Significant cortical activation was confirmed across the post-switch and mix phases (except for the left RLPFC (lRLPFC) in the post-switch phase), whereas no significant activation was observed in the pre-switch phase (Supplementary Table 3). Notably, activity was significantly greater in the right hemisphere compared to the left (t(1207) = 4.48, P < 0.001) and in the DLPFC compared to the RLPFC (t(1207) = 2.67, *P* = 0.008) (Fig. 3a,b,d). These results confirm that cognitive switching robustly recruits the LPFC, with a particular emphasis on the right and dorsal regions.

**Fig. 3.**
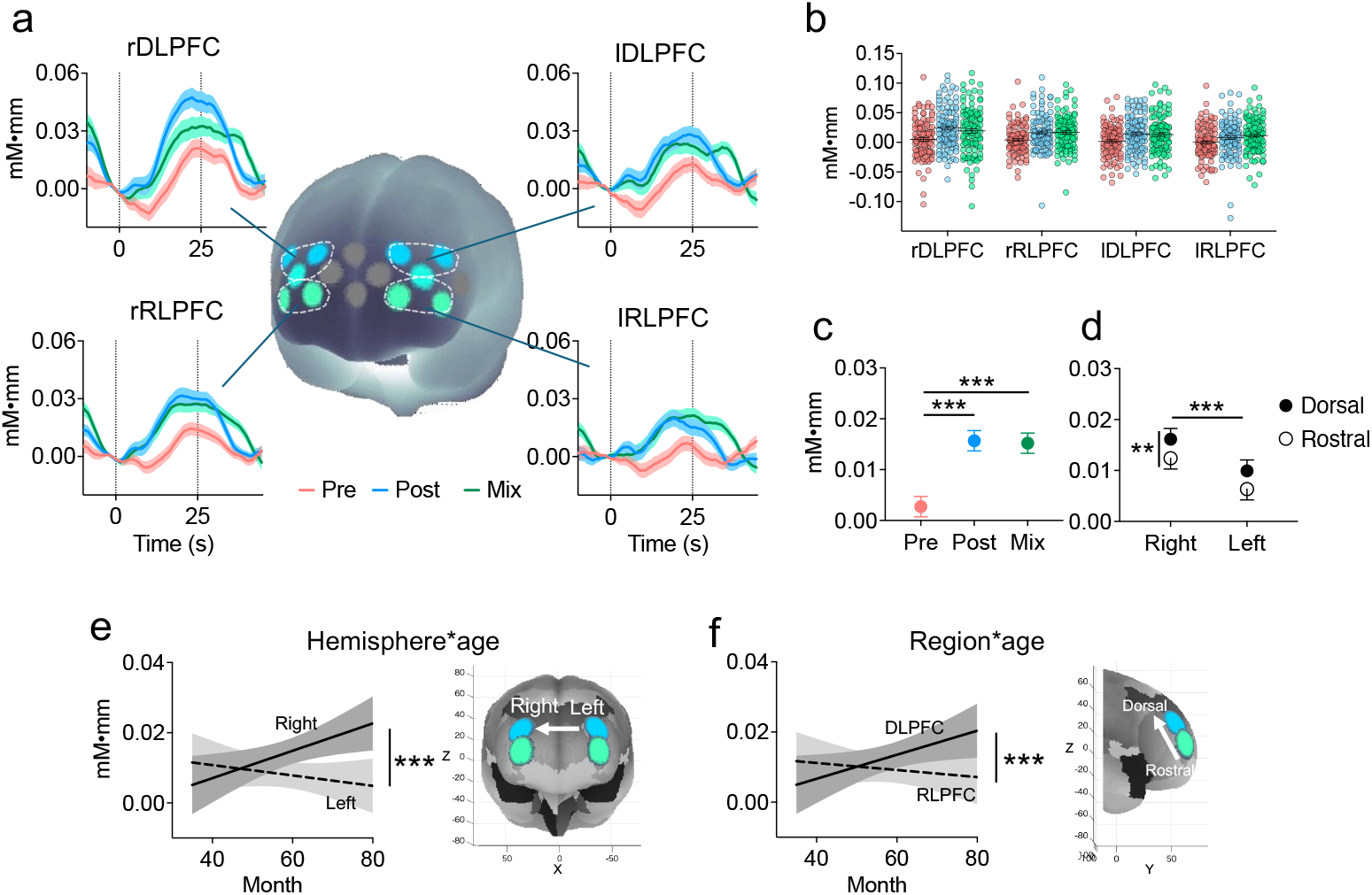
Task-related prefrontal activation and its modulation by age. **(a)** Grand-average waveforms (time courses) of oxy-Hb change for each task phase—pre (red), post (blue), and mix (green)—in the four regions of interest (ROIs: rDLPFC, lDLPFC, rRLPFC, lRLPFC). Shaded areas represent standard error (SE). Vertical dotted lines indicate onset (0 s) and offset (25 s) of the task block. **(b)** Distribution of mean oxy-Hb changes for each phase across all ROIs. Individual data points are shown. **(c, d)** Main effects on oxy-Hb signals (mM·mm) derived from the linear mixed-effects model (LMM). **(c)** Significant main effect of task phase: post (blue) and mix (green) > pre (red). **(d)** Significant main effects of hemisphere (right > left) and region (DLPFC > RLPFC). Error bars indicate SE. **(e, f)** Significant two-way interactions with age in months. Plots show the estimated marginal trends of oxy-Hb derived from the linear mixed-effects model. These illustrate that with increasing age, activation tends to increase in the right hemisphere and the dorsal region (DLPFC), while it tends to decrease in the post phase, the left hemisphere, and the rostral region (RLPFC). Shaded areas represent 95% confidence intervals. ***P < 0.001, **P < 0.01.

### Age-related changes in LPFC

To determine whether the prefrontal activation for cognitive switching was modulated by age in months, we compared the base model to a more comprehensive LMM that included age and all its interaction terms (Fig. 3e,f). A likelihood ratio test revealed that the model incorporating age provided a significantly better fit to the data (χ^2^ (12) = 61.92, *P* < 0.001), indicating that age is a critical factor in explaining the observed prefrontal activation. We therefore focused our subsequent analysis on the LMM. The ANOVA for the age-interaction model not only confirmed significant main effects of phase, hemisphere, and region but also revealed significant two-way interactions between age and phase (*F*(2, 1194.8) = 10.12, *P* < 0.001), hemisphere (*F*(1, 1195.8) = 22.15, *P* < 0.001), and region (*F*(1, 1196.1) = 15.03, *P* < 0.001). No higher-order interactions reached significance (all *Ps* > 0.19). To decompose these interactions, we performed post-hoc analyses on the estimated marginal trends. The hemisphere:age and region:age interactions converged to highlight a specific locus of age-related change. The effect of age on oxy-Hb signal change differed significantly between the hemispheres (*t*(1195) = 4.71, *P* < 0.001) (Fig. 3e) and between regions (*t*(1196) = 3.88, *P* < 0.001) (Fig. 3f). These effects were driven by a pattern wherein increasing age was associated with decreased activity in the left hemisphere and rostral subregion but increased activity in the right and dorsal subregions. This suggests a specific age-related functional differentiation in the PFC. In parallel, the phase:age interaction was driven by a significant difference in age-related trends between the mix and pre/post phases (pre vs mix: *t*(1193) = 3.33, *P* = 0.003; post vs mix: *t*(1195) = 4.28, *P* < 0.001). Taken together, these results demonstrate that the functional organization of this network during task switching undergoes significant reorganization across this period, which culminates in an increasing specialization to the rDLPFC.

### rDLPFC activity mediates relationship between sEBR and switching performance

Finally, we examined the relationship between PFC activation, sEBR, and task performance (Fig. 4). We conducted correlation analyses to test for specific associations between the activity of each of the four ROIs and our key measures: sEBR and switching accuracy. A Bonferroni correction was applied within each set of four comparisons (α = 0.05/4 = 0.0125). Only the activity of the rDLPFC was significantly and positively correlated with both sEBR (*rho* = 0.27, *P* = 0.004) (Fig. 4a) and switching accuracy (*rho* = 0.32, *P* < 0.001); no other ROIs showed significant correlations with either measure (*rho* < 0.20, *Ps* > 0.039; Supplementary Table 4). Additionally, we confirmed that the positive correlation between sEBR and rDLPFC activity was maintained after controlling for age (partial correlation: *rho* = 0.21, *P* = 0.027).

**Fig. 4.**
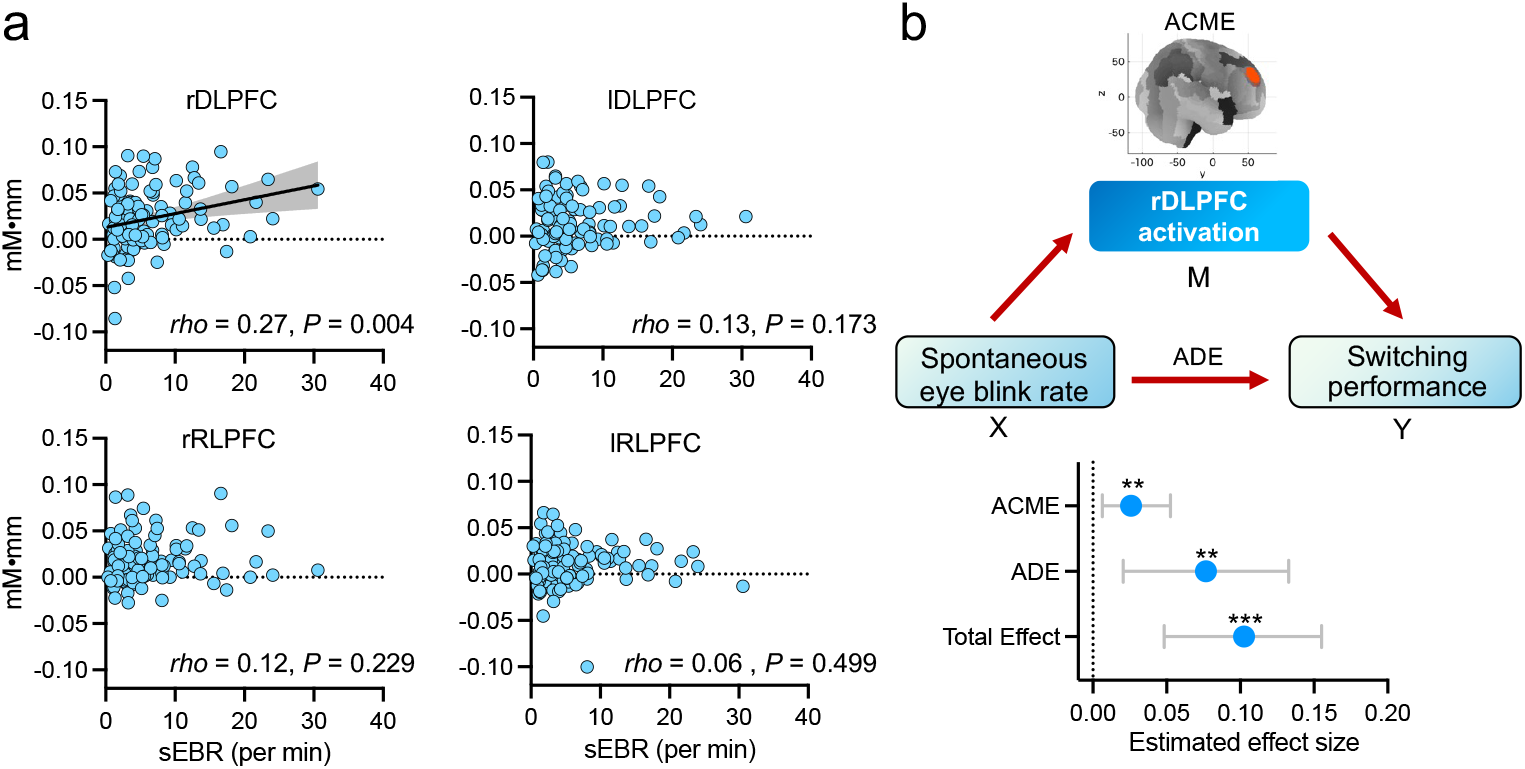
Activation of the right dorsolateral prefrontal cortex (rDLPFC) mediates the relationship between spontaneous eye-blink rate (sEBR) and switching performance. **(a)** Relationships between sEBR and PFC subregions. Only rDLPFC activation has a significant positive correlation with sEBR. **(b)** Top: A schematic diagram of the causal mediation model tested in this study illustrating the hypothesised pathway in which the effect of sEBR on switching performance is partially mediated by rDLPFC activation. Bottom: Estimated effect sizes from the causal mediation analysis. Points represent the effect estimates, and error bars represent 95% confidence. The total effect of sEBR on switching performance, the direct effect (average direct effect; ADE), and the indirect effect mediated through rDLPFC activation (average causal mediation effect; ACME) are shown. The confidence interval for both the ACME and ADE does not include 0, indicating that both the mediation effect and the direct effect are statistically significant. ***P < 0.001, **P < 0.01.

Next, to clarify how sEBR, rDLPFC activity, and switching accuracy relate to one another, we applied a cross-sectional mediation analysis (Fig. 4b) to decompose the sEBR–switching association into an indirect component via rDLPFC activity and a remaining direct component. The indirect component via rDLPFC was significant (Fig. 4b). Specifically, a shift in sEBR from the 25th to the 75th percentile was associated with a significant indirect effect on switching accuracy (Average Causal Mediation Effect (ACME) = 0.0258, 95% CI = [0.006, 0.0526], *P* = 0.005, estimated via quasi-Bayesian simulation). The mediated pathway accounted for approximately 24.5% of the total effect (95% CI = [5.71%, 62.65%]). These results provide support for our hypothesis, demonstrating that the rDLPFC recruitment is a significant pathway linking the neurodevelopmental processes indexed by sEBR to improvements in cognitive flexibility. The direct effect of sEBR on switching accuracy also remained significant (Average Direct Effect (ADE) = 0.0767, 95% CI = [0.0205, 0.1329], *P* = 0.010). This result suggests a partial mediation, indicating that this rDLPFC pathway is a candidate mechanism for the relationship between sEBR and switching performance. These associations were also robust to adjustment for sex and preschool (Supplementary Note 2).

## Discussion

The neural mechanisms driving the emergence of human higher-order cognition from infancy remain one of the most significant unsolved mysteries. Progress has been fundamentally constrained by our general inability to apply restrictive or invasive measurements to children, leaving the neural substrates of this critical developmental period poorly understood. Here, we provide the first evidence that emergent sEBR, a physiological output influenced by neuromodulatory systems, is associated with the development of executive function. Furthermore, the results of our fNIRS analyses identify the involvement of task-related functional differentiation in the PFC as an underlying neural basis of the link between sEBR and executive function development. Although there is a large amount of behaviour-based literature on executive function development^4,5,47^, this study demonstrates that the dramatic rise in sEBR during early childhood could serve as a trait-like physiological readout of the brain development required for complex cognition.

Our findings are the first to show that sEBR in early childhood is a neuro-behavioural marker predictive of switching performance. This is particularly relevant given the known developmental trajectory of sEBR, which is extremely low in newborns (approx. 0–5/min) and increases dramatically during development towards adult levels (approx. 20–30/min)^43–45^. Our data mirrored this pattern: sEBR was uniformly low with minimal individual variation in children under 4 years old, but significant individual differences emerged with age, with some children already blinking over 20 times/min (Fig. 2b). Importantly, the age-related emergence of inter-individual variability in sEBR allows sEBR to be used as a tool to index cognitive flexibility beyond chronological age, rather than relegating it to a metric that merely tracks age in parallel: sEBR predicted switching performance even after controlling for age in months. This age-adjusted association is consistent with inter-individual differences in executive-control circuit maturation in a developmental period when prefrontal recruitment is thought to become more selective and switching performance begins to diverge across children. In adults, inter-individual differences in sEBR have likewise been linked to executive function^32,48^, raising the possibility that early-emerging variability in sEBR reflects a trait-like signature that may persist into adulthood. This interpretation is consistent with evidence that individual differences in early executive function can forecast later outcomes in adults^1–3^, although longitudinal testing is required in order to verify this. While the neural origin for this blink increase is unclear, a leading candidate is the maturation of the brain dopaminergic system^49^. This hypothesis is supported by animal, clinical, and pharmacological evidence^24,27^. For instance, adults with Parkinson’s disease, characterised by nigrostriatal dopaminergic pathway dysfunction, exhibit both a markedly reduced sEBR—sometimes to levels comparable to young children^24,27,50^—and impaired executive function, especially task-switching^51,52^. Recent PET studies further confirm that sEBR in these patients correlates closely with symptom severity and dopamine depletion^28^. Therefore, the age-adjusted sEBR–switching association we have revealed in children is highly consistent with the adult and clinical literature, suggesting that sEBR in early childhood reflects the developing maturity of this dopaminergic system and frontostriatal network.

Despite the spatial constraints of fNIRS (restricted to the lateral prefrontal surface), we observed a clear age-related redistribution of task-evoked PFC activity. Specifically, activation became progressively more right-lateralised and dorsal-weighted across the preschool years, rather than changing uniformly across subregions (Fig. 3). This rightward and dorsal redistribution is consistent with the largely nonverbal DCCS, which often engages right-lateralised dorsal lateral frontoparietal control for task-set implementation and conflict resolution^7,19,23,53–55^. In contrast, left-lateralised regions linked to verbal processing and the more rostral LPFC implicated in exploratory/relational control^56,57^ may become less engaged as performance becomes more efficient. This result supports and extends previous reports identifying rDLPFC activation during the DCCS task^7,23,53,54^. Crucially, in a relatively large cohort spanning 35– 80 months, we show systematic shifts in task-evoked LPFC recruitment across the preschool years. This reorganization is consistent with accounts of functional specialization theory (diffuse-to-focal transitions; interactive specialization)^9–12^ and provides evidence that such specialization in the LPFC constitutes a neural basis for rapid early executive-function development.

It is plausible that the improvement of frontostriatal circuitry in functional efficiency promotes the functional differentiation of PFC recruitment. Our findings support this view: sEBR was specifically correlated with the recruitment of the rDLPFC, but not with any of the other prefrontal subregions measured (Fig. 4a). Strengthening this correlation, our mediation analysis identified a significant indirect effect, suggesting that sEBR reflects cognitive flexibility *via* the functional recruitment of the rDLPFC (Fig. 4b). This finding suggests that the link between sEBR and executive function development is underpinned by the functional maturation of the frontostriatal circuit, likely driven by neuromodulatory system development. This view aligns with the known rapid development of the dopaminergic system and our prior work linking dopamine-related genes (COMT) to both DCCS performance and rDLPFC activity in young children^23,58^.

Additionally, sEBR remained associated with switching performance even after accounting for rDLPFC activity. This suggests that sEBR may reflect components of executive function development not fully captured by LPFC activation measured with fNIRS. For example, developmental changes in broader frontostriatal coordination or subcortical contributions to rule updating could support switching with relatively efficient PFC recruitment^13,32^.

More broadly, although the mechanisms of sEBR development requires further validation, our findings position sEBR as a promising tool for assessing, and through which ultimately supporting, executive function in childhood. Executive function underpins a wide range of everyday activities, from following multistep instructions in the classroom to resolving social conflicts on the playground^2^. These capacities scaffold performance on complex cognitive tasks and behaviour regulation, providing a foundation for educational success and effective social interaction^1,59^. Thus, supporting executive function is a key educational priority. Importantly, guidance should be informed not only by behaviour but also by neural processes: measuring brain activity yields mechanistic insight that can improve the timing, targeting, and evaluation of interventions. In this context, sEBR is more than a convenient biomarker; it offers a non-invasive, low-cost, developmentally appropriate window into executive-function-related neurocognitive function that is otherwise difficult to access in early childhood. Because sEBR can be acquired by non-experts from ordinary digital video—and paired with AI-based automated detection^60^—it enables scalable at-home, preschool, and at-school data collection as well as “big-data” developmental science. Its minimal equipment requirements also make large-scale cross-cultural mapping of developmental trajectories feasible^58,61^, opening access to vast, previously untapped observations from diverse daily-life settings.

Several limitations should be noted, which also point to future directions for this line of exploration. First, the precise neuromodulatory mechanisms regulating sEBR remain an important open question. Blink generation involves multiple subcortical structures and brainstem areas^30,62–64^. It is likely a multifaceted process involving several neurotransmitter systems related to arousal, such as dopamine, acetylcholine, and noradrenaline^26,65^. Even within the well-studied dopaminergic system, the specific receptor interactions governing sEBR are not fully understood^26,40–42^. This complexity underscores the importance of future research into the fundamental mechanisms of blinking, not only to understand the behaviour itself, but also to fully comprehend its significance as a window into early brain development. Second, fNIRS signal interpretation remains debated because extracerebral/systemic physiology can contribute non-neural components^66^. We mitigated these concerns by restricting recordings to the frontal scalp (minimal hair interference) in a Japanese nursery-school cohort and by applying quality control with haemodynamic modality separation to reduce residual non-neural/systemic contamination. Finally, longitudinal work could lend additional support to the developmental trajectories and potential causal relationships implied by our findings.

In conclusion, this finding provides the first evidence in the 35-to-80-month age range that sEBR is associated with the development of executive function and the underlying prefrontal mechanisms. These findings indicate that blinking provides a unique and previously unrecognised insight into the neural development of young children.

## Methods

### Participants and procedures

Participants were children aged 2-6 years old recruited from two preschools in Osaka and Kyoto, Japan. All children were reported to have no developmental abnormalities, and prior informed consent was obtained from their parents. The study was conducted in accordance with the Declaration of Helsinki and was approved by the Ethics Committee of the Unit for Advanced Studies of the Human Mind, Kyoto University. Data from 113 children (55 female; mean age: 59 months, SD = 11.8, range: 35–80 months) were included in the final analyses. We were unable to obtain usable data from an additional 10 children due to experiment start refusal (n=6), refusal to wear the fNIRS cap (n=3), and task non-completion due to excessive movement (n=1).

### Sample size determination

Acknowledging that prior developmental neuroimaging studies have often faced limitations in statistical power due to small sample sizes^12^, the present study was designed a priori to obtain a large and robust sample. An a priori power analysis conducted with G*Power (v. 3.1) indicated that a sample of n = 82 was required to detect an anticipated medium-sized correlation (*r* = 0.30) with 80% power at a significance level of *α* = 0.05 (two-tailed). Our final analysed sample of n=113 fulfils this requirement, providing robust statistical power for our hypotheses.

### Behavioural task: Dimensional Change Card Sort (DCCS)

To measure executive function in young children, we used the DCCS task, a canonical measure of cognitive flexibility for this age^7,46^. We implemented a multi-stage version of the task suitable for a wide age range from early 3-year-olds to 6-year-olds, following the procedure from previous studies, including our own^23,53,54,67^. Stimulus cards consisted of two dimensions, “colour” and “shape”(Fig. 1a). In each trial, a target card was presented with two test cards, each matching the target on only one of the two dimensions. The task comprised three sessions, each using a different set of cards. Each session consisted of a pre-switch phase (colour rule), a post-switch phase (shape rule), and a mix phase (switching between both rules). Each sorting phase was limited to 25 s, during which participants attempted up to 8 trials. A 15-s rest period was provided between task phases. The order of rule presentation was fixed for all participants: “colour” for the pre-switch phase and “shape” for the post-switch phase. The mix phase followed a fixed sequence of rules: colour, shape, shape, colour, colour, shape, shape, colour. The primary outcome measure was the rule-switching success rate (%). This metric was defined as the accuracy on specific trials that required a rule switch, which is to say the first trial immediately following the transition from the pre-switch phase to the post-switch phase and up to 5 out of 8 trials within the mix phase for which the rule switched from that of the preceding trial, including the transition from the post-switch phase. The success rate was calculated as the percentage of correctly sorted trials out of the total number of these specific switch trials that the participant reached across all three sets.

### fNIRS recordings and analysis

We followed an fNIRS protocol refined through multiple studies by our group^7,23,53,54,68^, which has a confirmed methodological validity with DCCS in young children. Haemodynamic responses in the LPFC during the DCCS task were assessed using a multi-channel fNIRS optical topography system (OEG-16; Spectratech Inc., Tokyo, Japan) that emits near-infrared light at two wavelengths (770 nm and 840 nm). The configuration and placement of the fNIRS probe followed the procedures of our prior research^23,53,54^. The probe consisted of six light-source optodes and six detector optodes, forming 16 channels with a source–detector distance of 3 cm. The probe was placed with the lower row centred at Fpz. We performed spatial registration using an independent sample to project fNIRS channel locations into MNI standard space^69–71^. Probabilistic anatomical labeling^71^ provided anatomical reference^72^ for the a priori channel-based regions of interest (ROIs)^23,53,54,58^. The upper-channel ROI primarily covered the DLPFC (Brodmann area 9/46^72^), whereas the lower-channel ROI primarily covered the RLPFC (lateral Brodmann area 10, frontopolar area^72^) (Supplementary Table 1,2, Supplementary Note 3.). The acquired optical data were converted into changes in haemoglobin concentration based on the modified Beer-Lambert law. We did not assume a specific differential pathlength factor and calculated the relative change in oxygenated haemoglobin (oxy-Hb) in units of mM·mm^73^. Optical pathlength differences are reported to be minimal for symmetric hemispheric and adjacent regional comparisons^74^.

fNIRS data were analysed using proprietary software (OEG-16; Spectratech Inc.) and custom scripts in Python. Data were sampled every 0.655 s. Trials with visible motion contamination (body movement or probe displacement) were identified by synchronised video–fNIRS review and excluded; 98.4% of trials were retained. The data were smoothed using a 7-point moving average filter, following which, linear fitting was performed to reduce slow drifts.

Next, based on the haemodynamic-modality principle that task-evoked cortical activation typically produces an anti-correlated pattern between oxy-Hb and deoxy-Hb, whereas systemic physiological fluctuations, superficial perfusion, and some motion artifacts tend to induce positively correlated changes, we applied haemodynamic modality separation^75^ to each channel to decompose the signals into a functional component (putatively neural) and a systemic component (physiological noise/artifact) potentially capturing residual non-neural fluctuations related to subtle facial/motion activity (e.g., blinks). This approach has been described as a promising option for mitigating systemic contamination in fNIRS data^76^. Consistent with earlier paediatric applications, this procedure was robust in preschoolers and supported reliable estimation of DCCS-evoked responses^23,53,54,58^. Subsequent analyses used the functional component, which is expected to more closely reflect task-evoked cortical haemodynamics. We report oxy-Hb from the functional component, as oxy-Hb generally provides a higher signal-to-noise ratio than deoxy-Hb. To increase the signal-to-noise ratio, ROI signal values were calculated by averaging across channels (ch) for the rDLPFC (ch 2, 4, 5), lDLPFC (ch 11, 13, 14), rRLPFC (ch 3, 4, 6), and lRLPFC (ch 12, 13, 15), accounting for the similarity in optical properties of adjacent channels^74^. Channels spanning two regions (ch 4, 13) were included in the calculation for each region with a weight of 0.5 ^23,53,54,77^. Task-evoked activity was quantified as the mean oxy-Hb change in the task window (4.6 s post-onset to 20 s post-task) relative to the pre-onset baseline (-3.3 to 0 s), based on physiological plausibility and prior literature^32,54^. Supplementary robustness analyses indicated that the main results were not driven by this specific window choice (Supplementary Note 4; Supplementary Fig. 2). Finally, we applied an iterative Grubbs’ test (α = 0.0001) to the mean oxy-Hb values within each ROI to confirm that the results were not driven by rare artefactual extremes.

### Spontaneous eye-blink rate (sEBR)

sEBR was measured throughout the 6-min DCCS task (∼40% rest, ∼60% task). Each child’s face was recorded with a front-facing video camera (30 fps) positioned approximately 100–150 cm in front of the child. Two trained raters independently counted blinks for all videos (inter-rater correlation, *r* = 0.93), and their mean was used. A third rater independently scored a randomly selected 20% subset, showing high correlation with the average of the first two raters (*r* = 0.95). This video-based measurement has been used in prior studies; moreover, in adults it correlates well with vertical electro-oculography (vEOG), underscoring its validity^31,32^. Because attaching vEOG electrodes can impede task engagement and compliance in young children, the video-based approach provides the least intrusive and most feasible solution for preschooler assessments while maintaining measurement validity. For each participant, sEBR (blinks/min) was computed by dividing the total number of blinks during the 6-min recording by six (Fig. 1c). To minimize reactivity and encourage natural blinking, children were not explicitly told that blinking was the focus of the study. We adopted this DCCS-concurrent approach to obtain trait-like sEBR metrics; although dedicated resting-state blocks (e.g., cross-fixation) are used in adult studies^31,32^, they are pragmatically difficult for young children, and our design provides a valid, standardised alternative that improves cross-study comparability. Supplementary analysis confirmed the cross-context stability of the current method (Supplementary Fig. 2). Data were unavailable for one child due to insufficient eye visibility in the video, leaving the data for 112 participants for analyses.

### Analysis plan

Statistical analyses were primarily conducted in R (version 4.3.2), with supplementary analyses and data visualization performed in GraphPad Prism (version 9). The significance level was set at α = .05; corrections for multiple tests, when applied, are reported alongside the corresponding analyses. First, we assessed behavioral associations. Spearman’s rank correlations (*rho*) and partial correlations (controlling for age) were used to assess the relationships between age, sEBR, and switching accuracy. Second, we analysed oxy-Hb changes using linear mixed-effects models in R (using the “lme4”, “lmerTest”, and “emmeans” packages). We tested a comprehensive model including fixed factors of task phase (pre, post, mix), hemisphere (left, right), region (dorsolateral, rostrolateral), and age in months, along with all interaction terms, and participant as a random intercept. A likelihood ratio test confirmed this model’s superior fit against a base model without age terms. We used Type III ANOVA (Satterthwaite’s approximation) and post-hoc comparisons to analyse main effects and interactions. Third, to integrate brain and behaviour, we correlated the task switch-related activity (i.e., post and mix phases) of the four ROIs with sEBR and switching accuracy (Spearman’s *rho*) using a Bonferroni correction (α = .0125). Finally, we tested our proposed mechanism using causal mediation analysis (R package “mediation”) ^78,79^ to examine the pathway from sEBR (X) to switching accuracy (Y) mediated by rDLPFC activity (M). This analysis decomposes the total effect into the Average Causal Mediation Effect (ACME), representing the indirect effect via the mediator, and the Average Direct Effect (ADE), representing the direct effect not explained by the mediator. We assessed statistical significance using 95% confidence intervals from a quasi-Bayesian approximation^78,79^ (5,000 simulations). The sEBR effect was defined as the interquartile-range increase (2.13→8.04 blinks/min). Furthermore, a sensitivity analysis (medsens) was conducted to evaluate the robustness of the ACME to potential unmeasured confounding (Supplementary Note 1)^78^. We repeated the analyses with sex and preschool (site) as a covariate to evaluate whether sex influenced the results; the conclusions were unchanged (Supplementary Note 2).

## CRediT author statement

**Ryuta Kuwamizu**: Conceptualization, methodology, performing the experiments, data analysis, visualization, writing—original draft preparation, project administration, and funding acquisition; **Nozomi Yamamoto**: Performing the experiments, data analysis, and writing—review and editing; **Kota Otani**: Performing the experiments, data analysis, and writing—review and editing; and **Yusuke Moriguchi**: Conceptualization, writing—review and editing, supervision, and funding acquisition.

## Acknowledgements

The authors would like to thank their colleague Dr. Watanabe (Kyoto University) for their invaluable discussions. They thank Dr. Koizumi (University of Tsukuba, Japan) for blink analysis program. The authors express their gratitude to Ms. Noguchi (ELCS–English Language Consultation Services, Japan) for helping with the manuscript. Finally, we are deeply grateful to the staff of the participating kindergartens and nursery schools, as well as to all child participants and their guardians, for their generous cooperation. The illustration (girl in Fig. 1a) was drawn by Atsuko K.

