## Supplementary for "Emergent blink rate in early childhood is associated with neural origins of executive function"

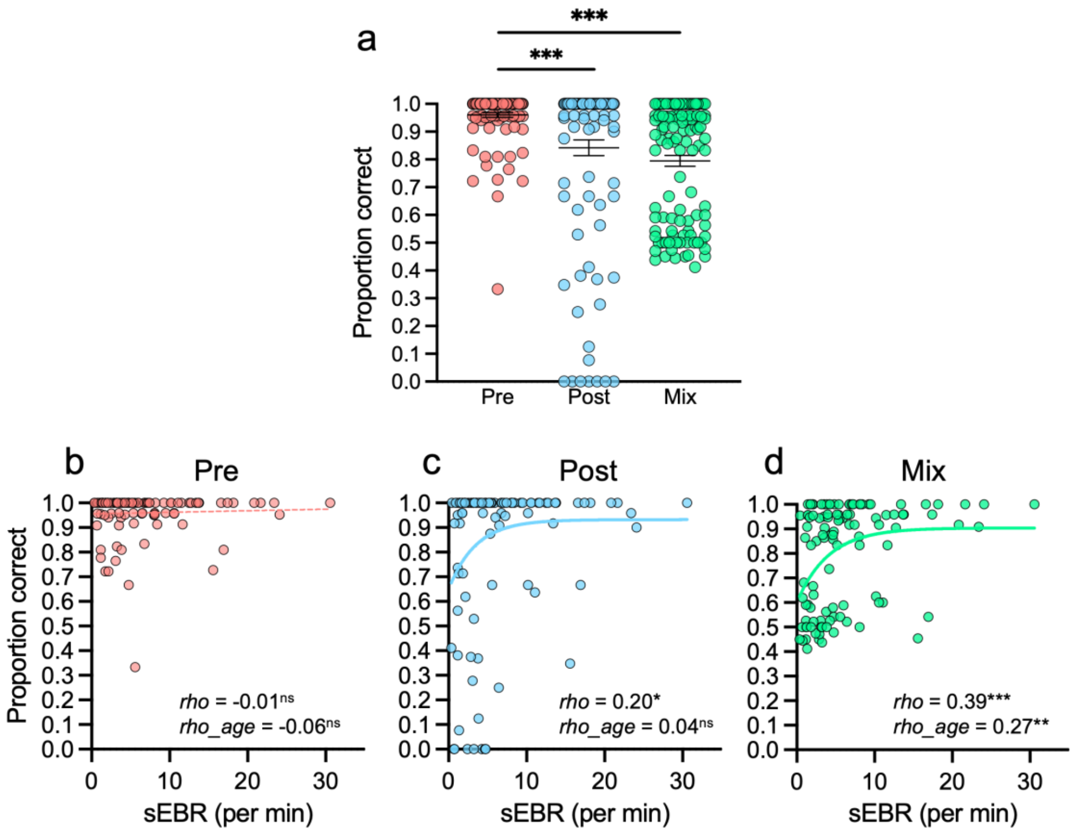


**Supplementary Fig. 1. Behavioral performance in the DCCS task and exploratory phase-wise associations with sEBR.**

**(a)** Accuracy in the three DCCS phases (Pre-switch, Post-switch, Mix). Each dot represents an individual child. The horizontal lines and error bars indicate the mean and standard error (SE), respectively. Consistent with previous research, correct responses on the DCCS task significantly decreased from the Pre-switch baseline phase to the more demanding Post-switch and Mix phases (Friedman test: $\chi^2(2) = 68.82$, $P < 0.001$; post-hoc Wilcoxon signed-rank tests with Bonferroni correction: both $P < 0.001$).

**(b–d)** Exploratory correlations between sEBR (blinks/min) and accuracy in the **(b)** Pre-switch, **(c)** Post-switch, and **(d)** Mix phases (Spearman's *rho*). The sEBR–accuracy association was most pronounced in the Mix phase (*rho* = 0.39, $P < 0.001$; partial Spearman's *rho* controlling for age = 0.27, $P = 0.005$). These exploratory analyses suggest that emergent sEBR relates preferentially to performance under higher cognitive flexibility demands (Mix phase), consistent with the main sEBR–switching performance results.

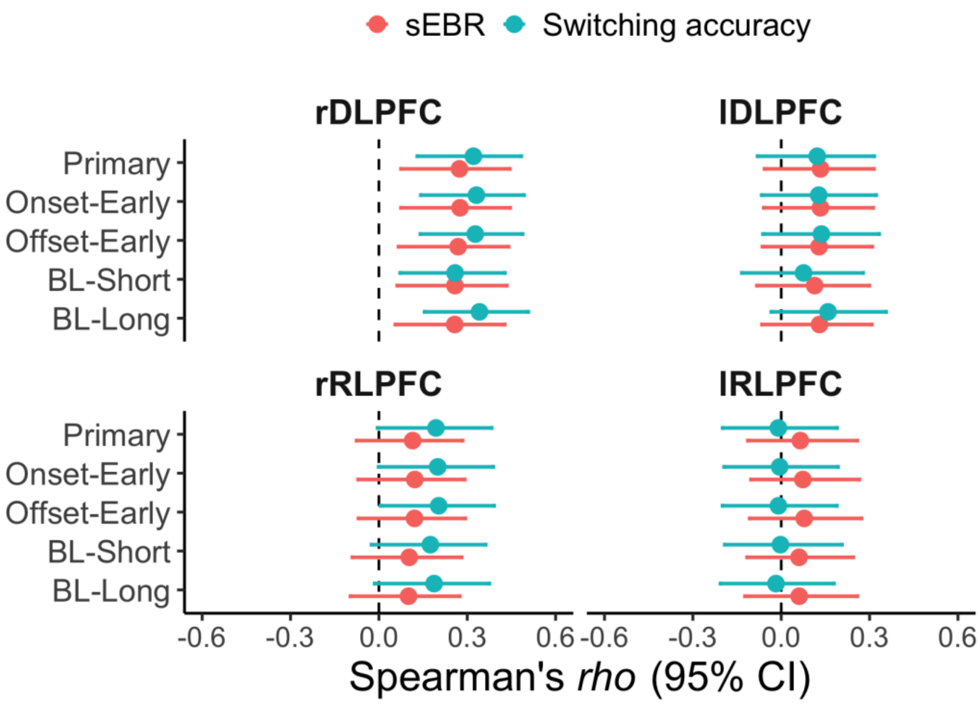


**Supplementary Fig. 2. Robustness of PFC ROI–behavior associations to window specification.**

Forest plots summarize Spearman correlations (*rho*) between PFC ROI responses and sEBR (red) and switching accuracy (blue) across alternative window specifications. The Primary specification had a baseline window from -3.3 s to 0 s and a post-onset averaging window from 4.6 s after onset to 20 s post-task (sampling interval ≈ 0.66 s). There were two baseline length variants, very short (BL-Short; from approximately -0.66 s to 0 s, ~1 sample) and longer (BL-Long; from approximately -5.2 s to 0 s, ~8 samples), each with an task window the same as that for the Primary specification. Task window variants had the same baseline window as for the Primary specification but the task window boundaries were shifted: For Onset-Early, the task window began immediately after task onset (from ≈0 s to 20 s post-task) and for Offset-Early the averaging window ended earlier than the Primary task window (from 4.6 s after onset to 13 s post-task). Points indicate correlation estimates and horizontal bars indicate 95% confidence intervals (CI).

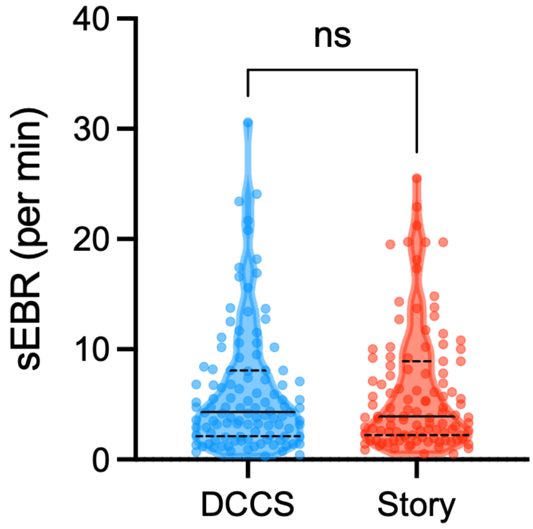


**Supplementary Fig. 3. Cross-context stability of spontaneous eye-blink rate (sEBR).**

To validate that our task-concurrent sEBR metric reflects a stable, trait-like physiological characteristic rather than a transient response to task demands, we compared sEBR measured during the DCCS task with that measured during a passive control condition (listening to narrated stories accompanied by illustrations[1]) in the participants with valid recordings for both conditions ($N$ = 110). The video recording and blink-counting procedures were identical to those used in the main task. A Wilcoxon signed-rank test revealed no significant difference in blink rates between the active DCCS task and the story-listening condition ($W$ = -83.00, $P$ = 0.90). The violin plots show the probability density of the data, with horizontal lines representing the median (solid) and quartiles (dashed). This result indicates that the executive demands of the DCCS did not systematically alter sEBR at the group level. Furthermore, a correlation analysis demonstrated a strong positive association between sEBR in the two conditions ($rho$ = 0.61, $P$ < 0.001).

**Supplementary Table 1. MNI coordinates for channels.**

Spatial registration was performed based on an independent sample of six children. The table reports estimated MNI coordinates (x, y, z; mean ± SD across participants) for channels used in the region of interest (ROI) definitions. See Supplementary Note 3 for details.

| Channel | MNI coordinates (mm) | | |
|---|---|---|---|
| | x | y | z |
| Ch2 | 45.5 ± 3.6 | 45.2 ± 3.4 | 28.7 ± 4.5 |
| Ch3 | 45.0 ± 3.8 | 57.5 ± 4.8 | 5.3 ± 5.9 |
| Ch4 | 37.0 ± 5.0 | 59.8 ± 3.3 | 19.3 ± 5.2 |
| Ch5 | 25.0 ± 4.8 | 59.5 ± 3.3 | 32.2 ± 5.3 |
| Ch6 | 25.2 ± 4.5 | 69.8 ± 1.8 | 10.3 ± 6.4 |
| Ch11 | -23.0 ± 4.0 | 57.8 ± 4.8 | 33.5 ± 4.3 |
| Ch12 | -25.5 ± 4.6 | 68.0 ± 3.3 | 10.0 ± 5.6 |
| Ch13 | -36.8 ± 5.0 | 57.2 ± 5.8 | 20.3 ± 4.6 |
| Ch14 | -44.5 ± 4.1 | 40.7 ± 7.8 | 30.7 ± 4.0 |
| Ch15 | -46.5 ± 4.2 | 51.5 ± 6.3 | 4.7 ± 5.5 |

**Supplementary Table 2. ROI probabilistic anatomical labels.**

Probabilistic anatomical labeling indicates the relative overlap probability of each region of interest (ROI) with anatomical regions (N = 6). ROI-level labeling distributions were computed using the same channel weights as used for ROI signal computation (boundary channels are shown in parentheses and were weighted as 0.5). **Abbreviations:** DLPFC, dorsolateral prefrontal cortex; RLPFC, rostrolateral prefrontal cortex; FPA, frontopolar area; OFA, orbitofrontal area; IFG, inferior frontal gyrus; r, right; l, left.

| ROI | Channel | Region (Brodmann area) [%] | Top label |
|---|---|---|---|
| rDLPFC | Ch2, (Ch4), Ch5 | DLPFC (9/46) [58.7%], Pars triangularis (45) [21.5%], FPA (10) [19.8%] | DLPFC (9/46) |
| lDLPFC | Ch11, (Ch13), Ch14 | DLPFC (9/46) [58.9%], Pars triangularis (45) [24.4%], FPA (10) [14.6%], Pars opercularis (44) [2.1%] | DLPFC (9/46) |
| rRLPFC | Ch3, (Ch4), Ch6 | FPA (10) [56.2%], DLPFC (9/46) [37.2%], OFA (11) [5.8%], Pars triangularis (45) [0.8%], IFG (47) [0.1%] | FPA (10) |
| lRLPFC | Ch12, (Ch13), Ch15 | FPA (10) [43.4%], DLPFC (9/46) [40.9%], Pars triangularis (45) [10.5%], OFA (11) [5.1%] | FPA (10) |

**Supplementary Table 3. One-sample t-test results for haemodynamic responses against baseline.** Statistical results of one-sample *t*-tests comparing the mean oxy-Hb change to zero (baseline) for each task phase and region of interest (ROI), corresponding to the data shown in Fig. 3b in the main manuscript. N denotes the number of participants. Asterisks indicate statistical significance after Bonferroni correction for multiple comparisons (12 tests: 3 phases × 4 ROIs; significance threshold α = 0.0042). (ns, not significant ($P$ >0.0042), ***$P$ < 0.000084).

| Phase | Region | Hemisphere | N | Oxy-Hb [mM·mm] | t-value | Cohen's d | $P_{uncorr.}$ | Sig. |
|---|---|---|---|---|---|---|---|---|
| Pre | DLPFC | Left | 107 | 0.002 | 0.79 | 0.08 | 0.434 | ns |
| | | Right | 112 | 0.005 | 1.68 | 0.16 | 0.096 | ns |
| | RLPFC | Left | 112 | 0.000 | -0.01 | 0.00 | 0.993 | ns |
| | | Right | 113 | 0.004 | 1.74 | 0.16 | 0.084 | ns |
| Post | DLPFC | Left | 106 | 0.014 | 4.94 | 0.48 | < 0.001 | *** |
| | | Right | 111 | 0.024 | 7.36 | 0.70 | < 0.001 | *** |
| | RLPFC | Left | 111 | 0.008 | 2.77 | 0.26 | 0.007 | ns |
| | | Right | 112 | 0.016 | 5.84 | 0.55 | < 0.001 | *** |
| Mix | DLPFC | Left | 107 | 0.013 | 4.24 | 0.41 | < 0.001 | *** |
| | | Right | 112 | 0.019 | 5.14 | 0.49 | < 0.001 | *** |
| | RLPFC | Left | 112 | 0.011 | 4.29 | 0.41 | < 0.001 | *** |
| | | Right | 113 | 0.017 | 6.39 | 0.60 | < 0.001 | *** |

**Supplementary Table 4. Spearman's rank correlations of functional brain activity with sEBR and switching accuracy.** The table presents Spearman's rank correlation coefficients (*rho*) and uncorrected *P*-values assessing the relationships between task-related haemodynamic responses in each region of interest (ROI), sEBR, and switching accuracy. N denotes the number of participants. Statistical significance was determined based on a Bonferroni-corrected threshold of α= 0.0125 (0.05/4).

| | sEBR | | | Switching accuracy | | |
|---|---|---|---|---|---|---|
| ROI | *N* | *rho* | $P_{uncorr.}$ | *N* | *rho* | $P_{uncorr.}$ |
| rDLPFC | 111 | 0.27 | 0.004 | 112 | 0.32 | <0.001 |
| lDLPFC | 106 | 0.13 | 0.173 | 107 | 0.12 | 0.210 |
| rRLPFC | 112 | 0.12 | 0.229 | 113 | 0.19 | 0.040 |
| lRLPFC | 111 | 0.06 | 0.499 | 112 | -0.01 | 0.918 |

### Supplementary Note 1. Sensitivity to unmeasured confounding (sequential ignorability)

We further assessed the robustness of the mediation effect against potential violations of the sequential ignorability assumption (i.e., the presence of unmeasured confounders). The analysis showed that the average causal mediation effect (ACME) would only be reduced to zero if the residual correlation (*rho*) reached approximately 0.3.

### Supplementary Note 2. Sex- and site-adjusted analyses

We tested the robustness of the key findings by adjusting for sex or measurement site (data were collected at two preschools in Osaka and Kyoto). Partial Spearman correlations remained significant after adjusting for sex (sEBR–rDLPFC: *rho* = 0.25, *P* = 0.008; sEBR–switching accuracy: *rho* = 0.30, *P* = 0.002; rDLPFC–switching accuracy: *rho* = 0.30, *P* = 0.001) and after adjusting for site (*rho* = 0.28, *P* = 0.003; *rho* = 0.34, *P* < 0.001; *rho* = 0.36, P < 0.001, respectively). In the developmental linear mixed-effects model of oxyHb, adding sex or site did not improve model fit (sex: $\chi^2(1) = 0.074$, *P* = 0.79; site: $\chi^2(1) = 0.19$, *P* = 0.66). Mediation (sEBR → rDLPFC activation → switching accuracy) remained significant when adjusting for sex (ACME = 0.0234, 95% CI [0.0044, 0.0500], *P* = 0.009; average direct effect (ADE) = 0.0580, 95% CI [0.0004, 0.1163], *P* = 0.050; proportion mediated = 27.9%, *P* = 0.014) and when adjusting for site (ACME = 0.0261, 95% CI [0.0067, 0.0525], *P* = 0.004; ADE = 0.0783, 95% CI [0.0243, 0.1327], *P* = 0.005; proportion mediated = 24.2%, *P* = 0.004). Overall, the central findings were robust to sex and site.

### Supplementary Note 3. Spatial registration and probabilistic anatomical labeling of fNIRS channels

We performed spatial registration in an independent sample of six children (2 female; mean age: 55 months, SD = 11.9, range: 42–72 months) recruited from the same age cohort to obtain an anatomical reference for the channel-based ROIs used in the main analyses. Head surfaces were captured using a Revopoint MIRACO Pro 3D scanner, and optode/channel locations were reconstructed with OEG-3DXYZ-OBJ (Spectratech, Japan). Channel positions were then registered to MNI standard space using NIRS-SPM[2], and probabilistic anatomical labeling was obtained using NFRI functions[3]. Brodmann area (BA) information was summarized with MRIcro[4].

### Supplementary Note 4. Robustness of window specification

To assess whether our main interpretation depends on a specific window choice, we confirmed the key PFC–sEBR analyses using a small set of physiologically plausible alternatives (Supplementary Fig. 2). The primary specification used a baseline window of -3.3 to 0 s and an averaging window of 4.6 s post-onset to 20 s post task. We then varied (i) baseline length while keeping the averaging window unchanged (very short baseline -0.66 to 0 s; long baseline -5.2 to 0 s), and (ii) averaging window boundaries while keeping the baseline window unchanged (Onset-Early from approximately 0 s to 20 s post-task, thus starting 4.6 s earlier; Offset-Early from 4.6 s post-onset to 13 s post-task, thus ending 6.6 s earlier). Across specifications, PFC–sEBR and PFC–switching accuracy associations showed similar point estimates with substantial overlap of the 95% confidence intervals, and ROI estimates were highly consistent across specifications (e.g., primary vs alternatives: *rho* = 0.961–0.998 for rDLPFC). These findings indicate that the main conclusions are robust to reasonable, physiology-motivated variations in window definition rather than being driven by a single boundary choice.
